# Omicron sublineage BA.2.75.2 exhibits extensive escape from neutralising antibodies

**DOI:** 10.1101/2022.09.16.508299

**Authors:** Daniel J. Sheward, Changil Kim, Julian Fischbach, Sandra Muschiol, Roy A. Ehling, Niklas K. Björkström, Gunilla B. Karlsson Hedestam, Sai T. Reddy, Jan Albert, Thomas P. Peacock, Ben Murrell

## Abstract

Several sublineages of omicron have emerged with additional mutations that may afford further antibody evasion. Here, we characterise the sensitivity of emerging omicron sublineages BA.2.75.2, BA.4.6, and BA.2.10.4 to antibody-mediated neutralisation, and identify extensive escape by BA.2.75.2. BA.2.75.2 was resistant to neutralisation by Evusheld (tixagevimab + cilgavimab), but remained sensitive to bebtelovimab. In recent serum samples from blood donors in Stockholm, Sweden, BA.2.75.2 was neutralised, on average, at titers approximately 6.5-times lower than BA.5, making BA.2.75.2 the most neutralisation resistant variant evaluated to date. These data raise concerns that BA.2.75.2 may effectively evade humoral immunity in the population.

## Main text

SARS-CoV-2 omicron sublineage BA.2.75 expanded rapidly in some parts of the world, but has so-far not outcompeted BA.5 globally. Despite similar geometric mean neutralising titers (GMT) to BA.5, BA.2.75 remained sensitive to classes of antibodies that BA.5 had escaped^1,2^, including potent cross-neutralising epitope group B and D1 antibodies^1^, suggesting significant scope for further antibody evasion. The emergence and rapid growth of a sublineage of BA.2.75 carrying additional mutations R346T, F486S, and D1199N (BA.2.75.2) (Fig 1A) suggested more extensive escape from neutralising antibodies.

**Fig. 1.**
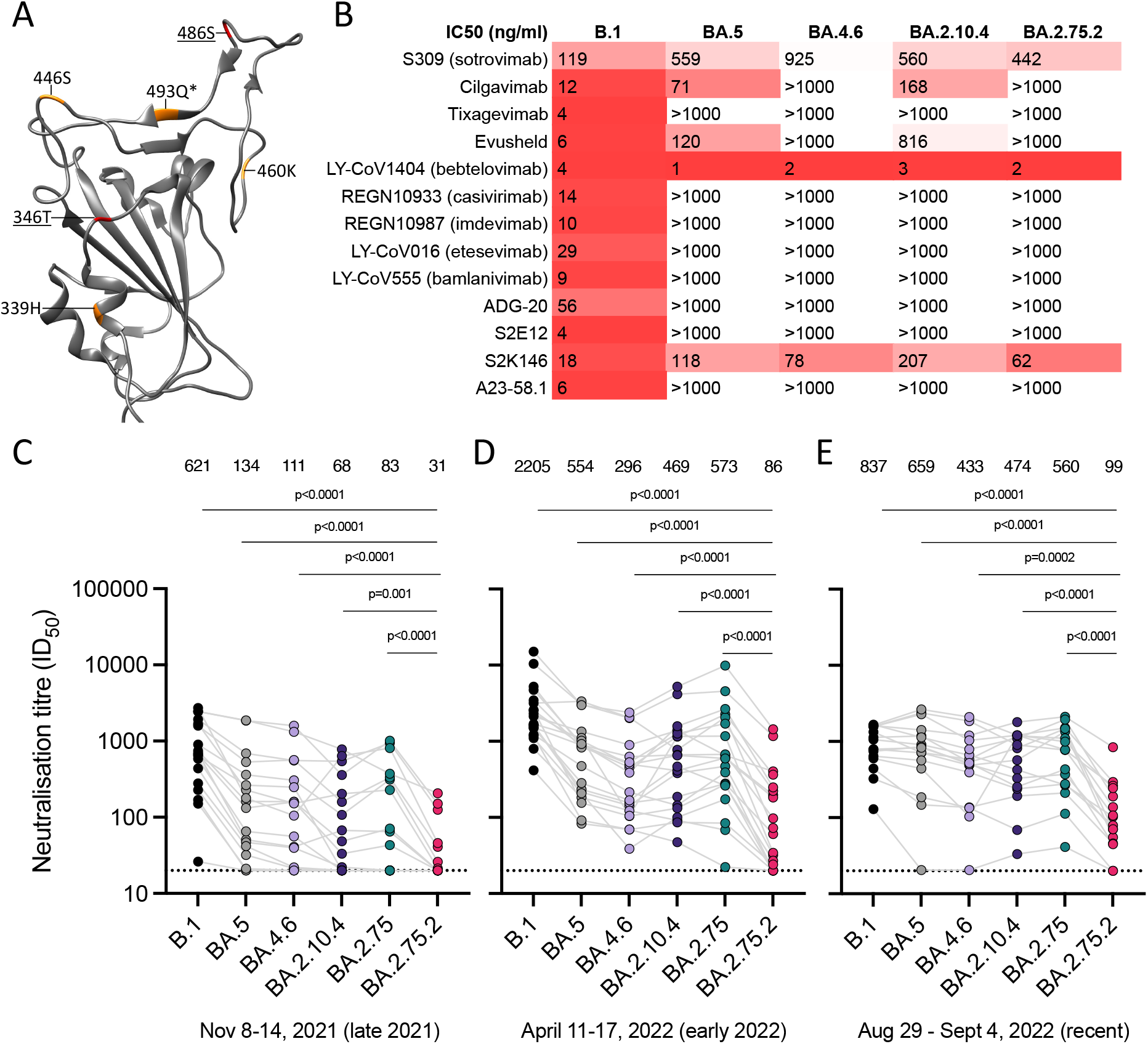
BA.2.75.2 escapes neutralising antibodies. **(A)** Differences from BA.2 in BA.2.75 (orange), and BA.2.75.2 (red, underlined), depicted upon the SARS-CoV-2 BA.2 RBD (pdb:7UB0). *indicates reversion. Sensitivity of SARS-CoV-2 omicron sublineages relative to B.1 (D614G) to neutralisation by **(B)** monoclonal antibodies, and randomly sampled sera from blood donated in Stockholm, Sweden between **(C)** 8-14 Nov 2021 (N=18), **(D)** 11-17 April 2022 (N=18) and **(E)** 29 Aug - 4 Sept 2022 (N=16). Sera with neutralisation <50% at the lowest dilution tested (20) are plotted as 20 (dotted line). ID_50_, 50% inhibitory dilution; IC_50_, 50% inhibitory concentration.

Mutations to other residues at these positions have been previously characterised: F486V mediates significant escape in BA.5^3^; R346K contributed to antibody escape in the pre-omicron Mu variant^4^, and occured in some early omicron BA.1 isolates^5^, and the widespread BA.1.1 lineage; and R346S occurred in a spike evolved *in vitro* to escape polyclonal antibody responses (PMS20)^6^. Spike residue 346 has been specifically highlighted for its capacity for additional escape in omicron^7^, and numerous sublineages of omicron have been detected carrying convergent mutations there. BA.4.6, carrying R346T and N658S, is currently the dominant 346T-carrying lineage, and has been detected across a wide geographic distribution. Similarly, several lineages are emerging carrying mutations at 486, including BA.2.10.4 harbouring a highly mutated spike including F486P.

Here we report the sensitivity of emerging omicron sublineages BA.2.75.2, BA.4.6, and BA.2.10.4 to neutralisation by a panel of clinically relevant and pre-clinical monoclonal antibodies, as well as by serum from blood donated in Stockholm, Sweden.

BA.2.75.2 and BA.4.6 both show complete escape from cilgavimab and the Evusheld combination, while BA.2.10.4 retains some sensitivity to cilgavimab (Fig 1B). S309 (sotrovimab) exhibits similarly low potency against BA.5, BA.2.75.2, and BA.2.10.4, with some further reduction against BA.4.6. Bebtelovimab still potently neutralises all variants tested.

In order to characterise the evolving resistance to immunity at the population level, we evaluated the sensitivity to neutralisation by serum from random blood donors in Stockholm, Sweden including: (i) A “late 2021” cohort, sampled prior to the emergence of Omicron (Nov 8th - 14th, 2021; Fig 1C), (ii) an “early 2022” cohort, sampled after a large wave of infections driven by BA.1 then BA.2 as well as the rollout of third vaccine doses (Apr 11-17, 2022; Fig 1D), and finally (iii) a “recent” cohort, (Aug 29 - Sept 4, 2022; Fig 1E) after the spread of BA.5.

In only the recent cohort, ancestral B.1 titers are at a similar level to those against omicron variants (Fig 1E), potentially as a result of omicron infections. Neutralisation of B.1 is no longer significantly different to BA.5 (p=0.5 for *recent,* vs p<0.0001 for *late 2021* and p<0.0001 for *early 2022).*

BA.4.6 and BA.2.10.4 are moderately more resistant to neutralisation than BA.5, with GMTs in the most recent samples of 433 (BA.4.6) and 474 (BA.2.10.4) compared to 659 (BA.5).

Across all three timepoints, neutralisation of BA.2.75.2 by serum antibodies was significantly lower than all other variants tested (Fig 1C-E). Both R346T and F486S mutations contribute to the significantly enhanced resistance of BA.2.75.2 compared to BA.2.75 (Fig S2). For samples from late 2021, prior to Omicron, the majority (11/18) had ID_50_ titers against BA.2.75.2 that were below the limit of detection (<20). In ‘post-Omicron’ samples (early 2022, and recent cohorts) BA.2.75.2 is neutralised with a GMT approximately 6.5-times lower than that of the currently-dominant BA.5, representing the most resistant variant characterised to date.

Taken together, these data identify profound antibody escape by the emerging omicron sublineage BA.2.75.2, suggesting that it effectively evades current humoral immunity in the population.

## Appendix

### Supplementary Figures

**Fig S1.**
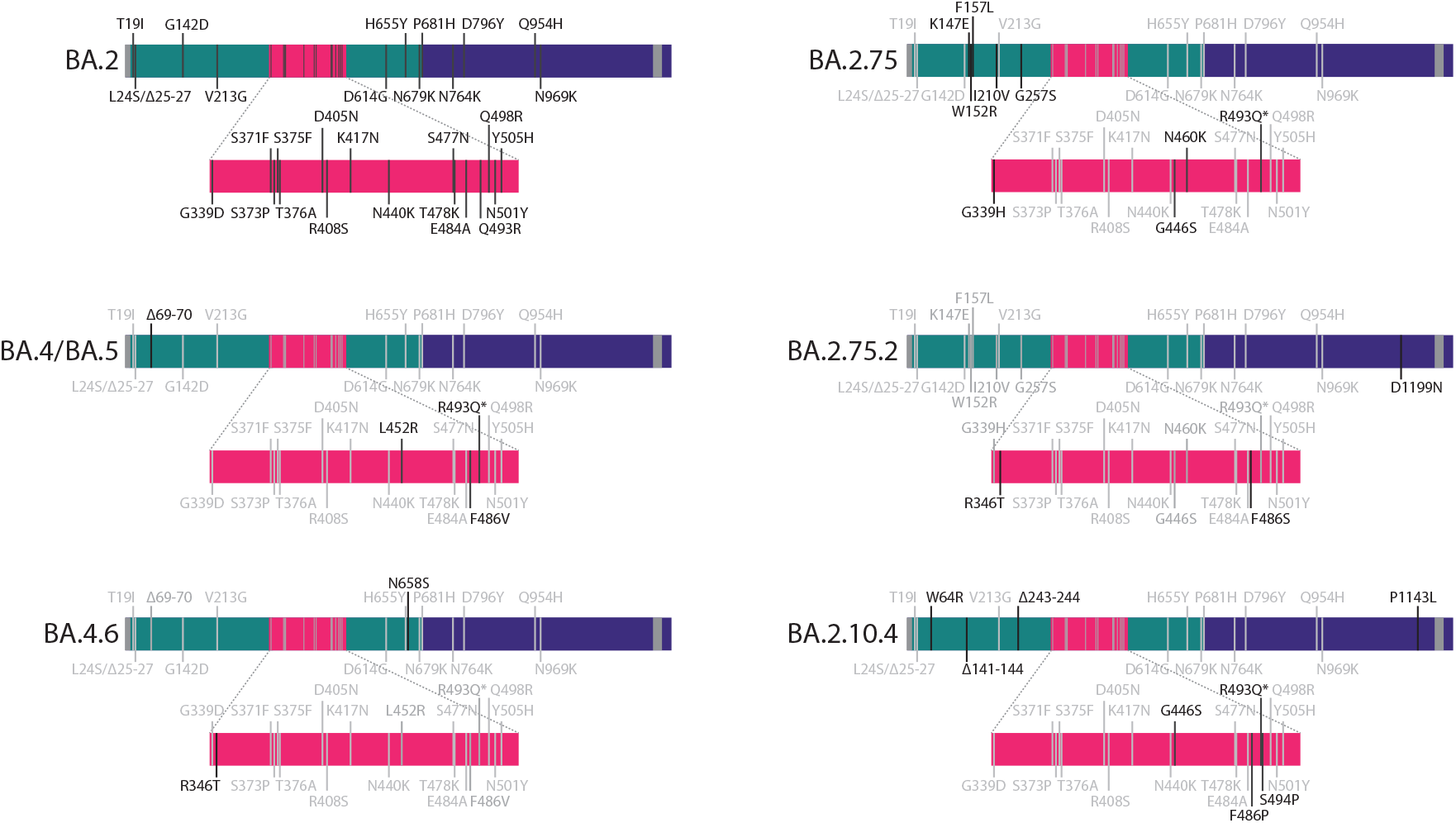
Spike mutations in omicron sublineages. Novel substitutions compared to the corresponding parental lineage are shown in black, with shared substitutions shown in grey. Depicted are BA.2, BA.4 and BA.5 (which have an identical spike amino acid sequence), BA.4.6^1^, BA.2.75^2^, BA.2.75.2^3^, and BA.2.10.4^4^.

**Fig S2.**
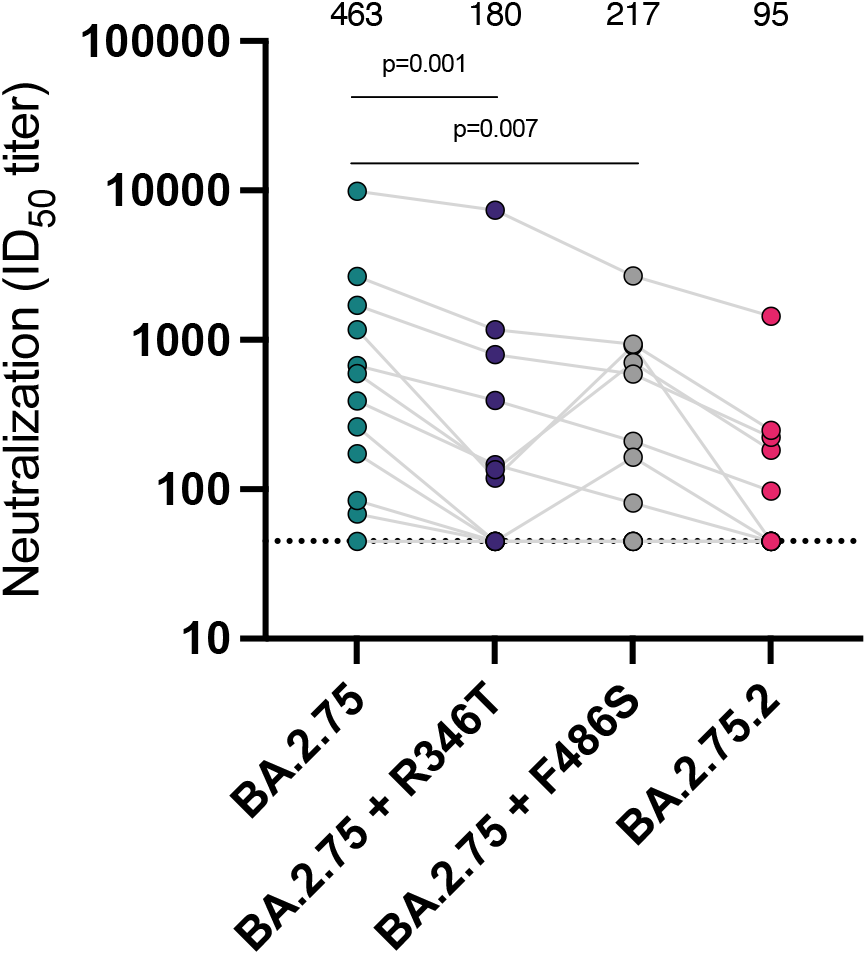
Both R346T and F486S contribute to BA.2.75.2 resistance. Neutralisation of BA.2.75 and mutants carrying changes present in BA.2.75.2 by serum samples (n=12) from 11-17 April 2022 highlights that both R346T and F486S contribute to the resistance of BA.2.75.2 relative to BA.2.75. Summarised above are the geometric mean neutralisation titers for each variant.

**Fig S3.**
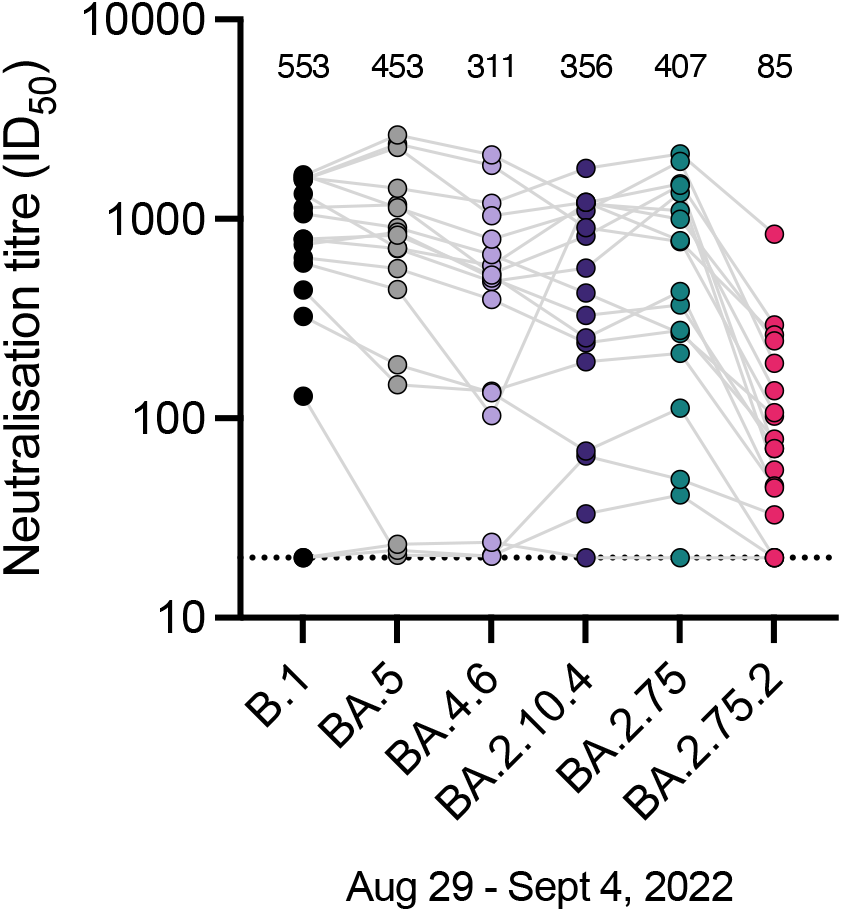
Neutralisation of omicron variants by serum from recently-donated blood: all samples. Sensitivity of SARS-CoV-2 omicron sublineages relative to B.1 (D614G) to neutralisation by randomly sampled sera from blood donated in Stockholm, Sweden between 29 August - 4 September 2022 (N=18). This includes samples that were excluded from Fig 1E due to neutralisation of B.1 <20 (plotted as 20, dotted line), for consistency with prior cohorts. ID_50_, 50% inhibitory dilution; IC_50_, 50% inhibitory concentration.

### Methods

#### Cell culture

HEK293T cells (ATCC CRL-3216) and HEK293T-ACE2 cells (stably expressing human ACE2) were cultured in Dulbecco’s Modified Eagle Medium (high glucose, with sodium pyruvate) supplemented with 10% fetal bovine serum, 100 units/ml Penicillin, and 100 μg/ml Streptomycin. Cultures were maintained in a humidified 37°C incubator (5% CO_2_).

#### Pseudovirus Neutralisation Assay

Pseudovirus neutralisation assay was performed as previously^5^. Briefly, spike-pseudotyped lentivirus particles were generated by co-transfection of HEK293T cells with a relevant spike plasmid, an HIV gag-pol packaging plasmid (Addgene #8455), and a lentiviral transfer plasmid encoding firefly luciferase (Addgene #170674) using polyethylenimine. Spike variants were generated by multi-site directed mutagenesis of BA.2, BA.4, or BA.2.75 expression plasmids^6^, and all mutated plasmids were subsequently confirmed by Sanger sequencing.

Neutralisation was assessed in HEK293T-ACE2 cells. Pseudoviruses titrated to produce approximately 100,000 RLU were incubated with serial 3-fold dilutions of serum or monoclonal antibody for 60 minutes at 37°C in a black-walled 96-well plate. 10,000 HEK293T-ACE2 cells were then added to each well, and plates were incubated at 37°C for 44-48 hours. Luminescence was measured using Bright-Glo (Promega) on a GloMax Navigator Luminometer (Promega). Neutralisation was calculated relative to the average of 8 control wells infected in the absence of antibody. Samples were run against all variants ‘head-to-head’ using the same dilutions.

#### Monoclonal antibodies

Cilgavimab and tixagevimab were evaluated as their clinical formulations. For the rest of the monoclonal antibodies evaluated, antibody sequences were extracted from deposited RCSB entries, synthesised as gene fragments, cloned into pTWIST transient expression vectors by Gibson assembly or restriction cloning, expressed and purified, all as previously described^7^.

#### Serum samples

Serum samples from anonymized blood donors from Stockholm, Sweden, were obtained from week 45, 2021 (8-14 November 2021) (prior to the BA.1/BA.2 Omicron infection wave), from week 15, 2022 (11-17 April 2022) (after the BA.1/BA.2 Omicron infection wave, but prior to the arrival of BA.4 or BA.5), and from week 35, 2022 (29 August - 4 September 2022) (after the spread of BA.5). Samples from the late 2021 and early 2022 cohorts were a subset of samples previously studied^5^, which had excluded samples that did not neutralise ancestral B.1, prior to assaying neutralisation across all variants. To ensure comparability with this previous sampling, the *recent* cohort in the main figure and analysis uses the same inclusion criterion, but for completeness we include the full set of neutralisation data in the SI (Fig S3). Sera were heat inactivated at 56°C for 60 minutes prior to use in neutralisation assays.

#### Ethical Statement

The blood donor samples were anonymized, and not subject to ethical approvals, as per advisory statement 2020–01807 from the Swedish Ethical Review Authority.

#### Statistical analysis

Individual ID_50_ and IC_50_ values for each sample against each variant were calculated in Prism v9 (GraphPad Software) by fitting a four-parameter logistic curve to neutralisation by serial 3-fold dilutions of serum/antibody. Neutralising titers between groups were compared using a Wilcoxon matched-pairs signed rank test using Prism v9.

## Author contributions

Conceptualization, D.J.S., T.P.P., B.M.;

Formal analysis, D.J.S.;

Conducted the assays, D.J.S., C.K., J.F., T.P.P.;

Designed the methodology, D.J.S., C.K., R.E., T.P.P, B.M.;

Responsible for figures and tables, D.J.S., T.P.P., B.M.;

Resources, S.M., R.E., S.R, N.K.B., G.B.K.H., J.A., B.M.;

Oversaw the study, D.J.S., G.B.K.H., S.T.R., J.A., B.M;

Funding Acquisition, S.T.R., G.B.K.H., J.A., T.P.P., B.M.

Writing – original draft, D.J.S., B.M.;

Writing – review & editing, D.J.S., G.B.K.H., J.A., T.P.P., B.M.

D.J.S and B.M. were responsible for the decision to submit the manuscript for publication.

## Acknowledgements

pCMV-dR8.2 dvpr was a gift from Bob Weinberg (Addgene plasmid # 8455; http://n2t.net/addgene:8455; RRID:Addgene_8455). pBOBI-FLuc was a gift from David Nemazee (Addgene plasmid # 170674; http://n2t.net/addgene:170674; RRID:Addgene_170674). We also wish to thank the clinicians and researchers, globally, who contributed to the sampling, sequencing, and identification of emerging SARS-CoV-2 variants.

## References

1 Cao Y, Yu Y, Song W, et al. Neutralizing antibody evasion and receptor binding features of SARS-CoV-2 Omicron BA.2.75. bioRxiv. 2022;: 2022.07.18.500332.

2 Sheward DJ, Kim C, Fischbach J, et al. Evasion of neutralising antibodies by omicron sublineage BA.2.75. Lancet Infect Dis 2022; published online Sept 1. DOI:10.1016/S1473-3099(22)00524-2.

3 Cao Y, Yisimayi A, Jian F, et al. BA.2.12.1, BA.4 and BA.5 escape antibodies elicited by Omicron infection. Nature 2022; 608: 593–602.

4 Halfmann PJ, Kuroda M, Armbrust T, et al. Characterization of the SARS-CoV-2 B.1.621 (Mu) variant. Sci Transl Med 2022; 14: eabm4908.

5 Cele S, Jackson L, Khoury DS, et al. Omicron extensively but incompletely escapes Pfizer BNT162b2 neutralization. Nature 2021; 602: 654–6.

6 Schmidt F, Weisblum Y, Rutkowska M, et al. High genetic barrier to SARS-CoV-2 polyclonal neutralizing antibody escape. Nature 2021; 600: 512–6.

7 Greaney AJ, Starr TN, Bloom JD. An antibody-escape estimator for mutations to the SARS-CoV-2 receptor-binding domain. Virus Evol 2022; 8: veac021.

## Supplementary References

1. pango-designation. Github https://github.com/cov-lineages/pango-designation/issues/741 (accessed Sept 16, 2022).

2. pango-designation. Github https://github.com/cov-lineages/pango-designation/issues/773 (accessed Sept 16, 2022).

3. pango-designation. Github https://github.com/cov-lineages/pango-designation/issues/963 (accessed Sept 16, 2022).

4. pango-designation. Github https://github.com/cov-lineages/pango-designation/issues/898 (accessed Sept 16, 2022).

5. Sheward DJ, Mandolesi M, Urgard E, et al. Beta RBD boost broadens antibody-mediated protection against SARS-CoV-2 variants in animal models. Cell Rep Med 2021; 2: 100450.

6. Sheward DJ, Kim C, Fischbach J, et al. Evasion of neutralising antibodies by omicron sublineage BA.2.75. Lancet Infect Dis 2022; published online Sept 1. DOI:10.1016/S1473-3099(22)00524-2.

7. Sheward DJ, Kim C, Ehling RA, et al. Neutralisation sensitivity of the SARS-CoV-2 omicron (B.1.1.529) variant: a cross-sectional study. Lancet Infect Dis 2022; 22: 813–20.

